# The most common RNF43 mutant G659Vfs41 is fully functional in inhibiting Wnt signaling and unlikely to play a role in tumorigenesis

**DOI:** 10.1101/711382

**Authors:** Jianghua Tu, Soohyun Park, Wangsheng Yu, Sheng Zhang, Ling Wu, Kendra Carmon, Qingyun J. Liu

**Affiliations:** Texas Therapeutics Institute and The Brown Foundation Institute of Molecular Medicine, The University of Texas Health Science Center at Houston, 1825 Pressler St., Houston Texas, USA

## Abstract

RNF43 is an E3 ligase that inhibits Wnt signaling by ubiquitinating Wnt receptors for degradation. It is mutated in various cancer types with the most recurrent mutation being the frameshift G659Vfs41 with frequencies of ~5-8% in colon, stomach and endometrial cancers. The mutation has always been assumed to be loss-of-function that would lead to increased Wnt signaling and tumorigenesis, yet no functional characterization has been reported. We analyzed the distribution of the G659Vfs41 mutation and its association with other cancer gene mutations, and found that the mutation occurred nearly exclusively in tumors with low expression of the DNA mismatch repair gene MLH1. Mutant RNF43-G659Vfs41 was no different from wild type RNF43 in expression, localization, R-spondin binding, promotion of Wnt receptor degradation, inhibition of Wnt signaling. We also found that colon tumors with RNF43-G659Vfs41 had low expression of Wnt/β-catenin signaling markers and were frequently mutated in BRAF. A colon cancer cell line with RNF43-G659Vfs41 and BRAF-V600E mutations was uniquely sensitively to activation of Wnt/β-catenin signaling. These findings indicate that RNF43-G659Vfs41 is likely a passenger mutation in MLH1-down regulated tumors, not a loss-of-function driver mutation that facilitates tumorigenesis. Tumors with RNF43-G659Vfs41 and BRAF-V600E may be susceptible to activators of Wnt signaling.

## Introduction

RNF43, a member of the RING finger E3 ubiquitin ligase family, was originally identified as an oncoprotein with upregulated expression in colon cancer^1^. It is a type I integral membrane protein with an extracellular domain of 175 amino acid (AA) residues, a single transmembrane domain, and an intracellular region of 565-AA with the RING finger E3 ligase domain (AA270-316) immediately following the transmembrane domain. A series of subsequent studies revealed that RNF43 and its related paralog ZNRF3 ubiquitinate the frizzled (FZD) family of Wnt receptors for degradation and thus serve as negative regulator of Wnt signaling^2–5^. Meanwhile, genome-wide sequencing of tumor samples uncovered that RNF43 and ZNRF3 were mutated in a wide variety of tumor types, with relatively high frequency (near 10%) found in pancreatic^6^, uterine endometrial^7^, stomach^8^, and colon cancers^7,9–14^. In the COSMIC cancer database, approximately half of the 1,036 mutations of RNF43 identified from all cancer types are either non-sense or frameshift mutations^15^. Some of the recurrent non-sense mutations of RNF43, such as R132x and R145x, are expected to produce non-functional proteins due to the lack of the E3 ligase domain, as well as missense mutations in the RING domain were shown to cause loss-of-function^4,16^. In pancreatic cancer, RNF43 mutations located inside or upstream the E3 ligase domain were identified and cell lines with such mutations were shown to be sensitive to inhibition of Wnt ligand secretion^17^. In colon cancer, RNF43 mutations were largely exclusive with APC mutation^7^. All these observations led to a seemingly obvious conclusion that nonsense and frameshift RNF43 mutations are loss-of-function mutations that would lead to tumorigenesis in a significant subset of gastrointestinal and endometrial cancers due to increased Wnt signaling^7,14^. Inhibition of Wnt ligand production is purported to provide an effective treatment for tumors with RNF43 non-sense or frameshift mutations^7,14^.

Remarkably, the single most common recurrent mutation of RNF43 is the deletion of a G in a seven G repeat near the 3’ end of its open reading frame (nucleotide 1969-1976), accounting for approximately half of all RNF43 mutations detected in colon, stomach, and endometrial cancer with an overall frequency of 5-8%^7,14,15^. This mutation, designated G659Vfs41, results in the truncation of the enzyme at Gly659 and shifts the reading frame to add a neopolypeptide of 41-AA (Fig. 1A)^15^. The mutation removes the C-terminal 123-AA region that is not essential for down regulation of Wnt signaling as it is far downstream of the E3 ubiquitin ligase domain^4,5^. Nevertheless, the mutation has always been proposed to be loss-of-function and play a role in colon cancer formation/progression due to disinhibition of Wnt signaling^7,14^. If and how RNF43-G659Vfs41 will affect its enzyme activity in regulation of Wnt signaling has never been reported. Both RNF43 and ZNRF3 ubiquinate lysine residues of FZD receptors for endocytosis and degradation^2–5^. The process is suggested to occur following the activation of Wnt signaling in which the E3 ligases interact with the disheveled (DVL) protein bound to the Wnt signaling complex to catalyze ubiquitination of FZD^4,5^. However, ubiquitination and degradation of FZD was shown to happen without concomitant activation of Wnt signaling, at least with recombinant enzymes and receptors^4^. E3 ubiquitin ligase activity of RNF43 and ZNRF3 can be inhibited by R-spondins, a group of four related secreted proteins (RSPO1-4) that strongly potentiate Wnt signaling and drive tumorigenesis when over-expressed by gene fusion or other mechanisms^18,19^. Here we show that the RNF43-G659Vfs41 mutant is actually fully functional in promoting FZD degradation and inhibition of Wnt signaling. In the three cancer types with highest incidence of G659Vfs41, nearly all tumors with this mutation had low expression of MLH1, a major player of DNA mismatch repair.

**Figure 1.**
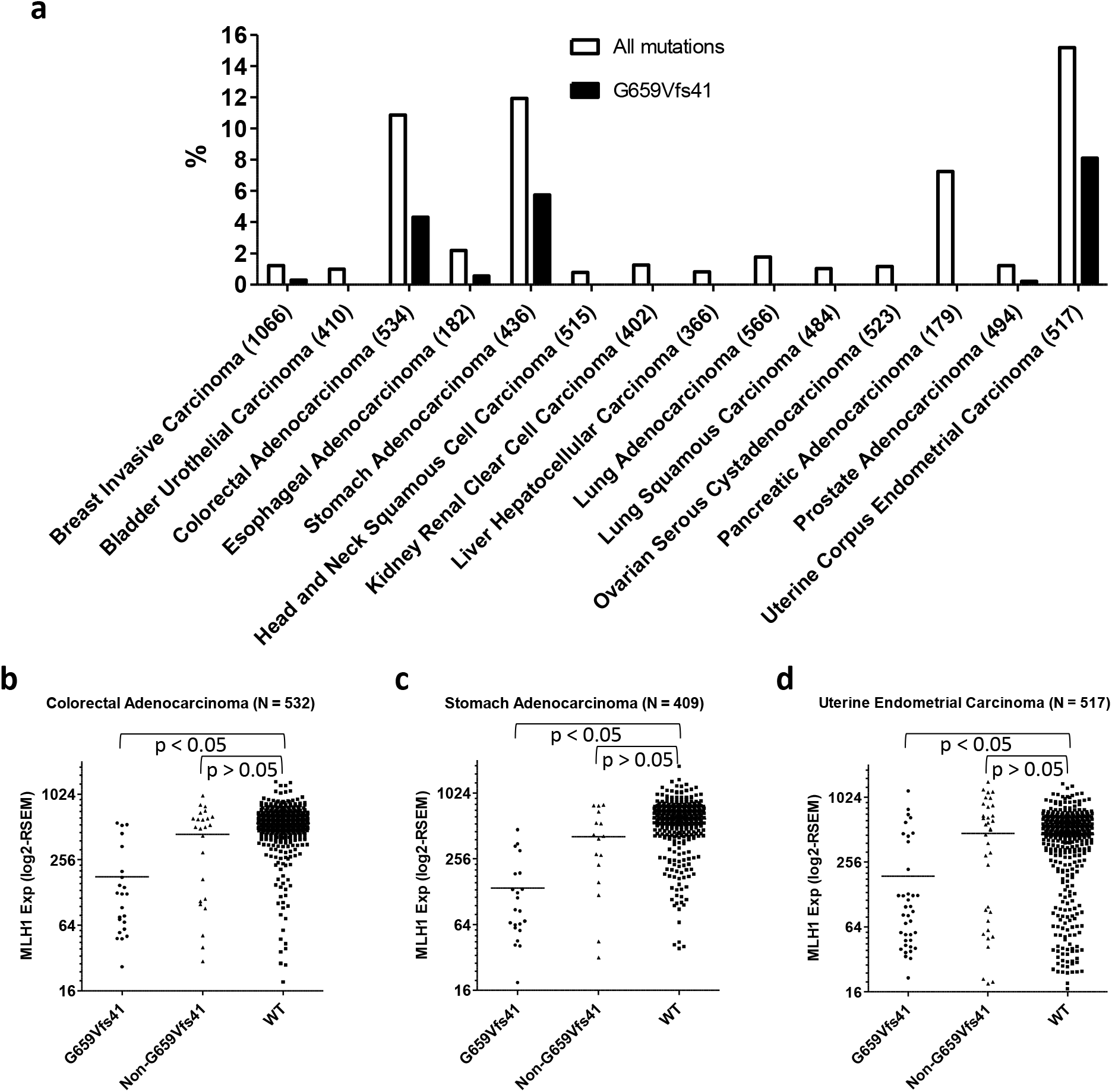
RNF43-G659Vfs41 is most common in colon, stomach, and endometrial cancers and the mutation is strongly associated with low MLH1 expression. a, Distribution of RNF43 mutations in all major types of solid tumors. b-d, Scatter plot of xpression of MLH1 in relation to mutation status of RNF43 in colorectal cancer (b), stomach cancer (c) and endometrial cancer (d). p values were calculated using one-way ANOVA followed by Dunn’s multiple comparison test.

Furthermore, in colon cancer, RNF43-G659Vfs41 mutations are most frequently associated with BRAF-V600E mutation and low Wnt/β-catenin signaling. Intriguingly, we show that a colon cancer cell line with RNF43-G659Vfs41 and BRAF-V600E mutations is highly sensitive to activation of Wnt signaling in vitro, resulting in suppression of cell growth and apoptosis. This suggests that tumors with BRAF-V600E mutation may require low Wnt signaling to survive. Overall, these findings indicate that RNF43-G659Vfs41 is not a driver in colon cancer or other cancer types and that colon tumors with BRAF-V600E mutation may be treated with activators of Wnt signaling.

## Results

### RNF43-G659Vfs41 is found nearly exclusively in cancers with low MLH1 expression

We first systematically analyzed RNF43 mutations identified in all major types of solid tumors in the TCGA database using cBioPortal^20,21^. Colon, endometrial, pancreatic, and stomach cancers showed the highest incidences of all RNF43 mutations with frequencies of 10.9%, 15.2%, 7.3%, and 11.9%, respectively (Fig. 1a). In colon, endometrial, and stomach cancers, G659Vfs41 accounted for 40% (23/81), 53% (42/79), and 48% (25/77) of all RNF43 mutations, respectively, whereas not a single G659Vfs41 mutation (0/13) was found in pancreatic cancer (p <0.05 vs any of the other three cancer types, Fisher’s exact test). The only other cancer type with G659Vfs41 mutation being identified more than once is breast cancer (3/16 RNF43 mutations in 1066 cases). These data strongly suggest that RNF43-G659Vfs41 occurs nearly exclusively in colon cancers, endometrial, and stomach cancers.

Previously, it was reported that RNF43 mutations are frequently associated with high microsatellite instability (MSI-H) in colon and stomach cancer^7,11,14^. We examined the relationship between RNF43 mutations and MLH1 expression in this TCGA cohort as low expression of MLH1 due to methylation is the most common cause of MSI-H ^22^. In colon cancer, RNF43-G659Vfs41 tumors showed a much lower expression of MLH1 when compared to those of RNF43-WT tumor (mean RNA-seq RSEM value of 179 vs 553, p <0.05) (Fig. 1b). In contrast, tumors with non-G659Vfs41 mutations in RNF43 showed no significant difference with those of WT in MLH1 expression (mean RNA-seq RSEM value of 440 vs 553, p > 0.05) (Fig. 1b). Similar data were found in endometrial and stomach cancers (Fig. 1c-d, Supplementary Fig. S2a-c). In pancreatic cancer in which RNF43 was mutated at relative high frequency except no G659Vfs41 mutation was identified, none of the tumors had low MLH1 expression (Supplementary Fig. S2d). Furthermore, of all solid tumors, only colon, stomach, and uterine endometrial cancers have a significant subset of tumors with low MLH1 expression (http://firebrowse.org/viewGene.html?gene=mlh1). Taken together, these results unequivocally show that RNF43-G659Vfs41 mutation in colon, endometrial and stomach cancers primarily occurred in tumors with low MLH1 expression and presumably defective DNA mismatch repair, suggesting that the 7 G repeat (bp1969-1976) of the RNF43 coding sequence is prone to single bp deletion in the absence of proficient DNA repair.

### No discernable difference between RNF43-G659Vfs41 and wild type in cellular localization and RSPO-binding

To examine functional consequences of the G659Vfs41 mutation, we generated constructs expressing the mature forms of full-length wild type (WT) RNF43 or G659Vfs41 with an HA tag (Fig. 2a). An additional construct with a termination codon immediately following G659 (G659x) was also generated to determine if the 41-AA neopolypeptide affects expression. The three constructs were transfected into HEK293T cells and Western blot (WB) analysis showed that they were expressed at similar levels with the expected differences in molecular weights (Fig. 2b), indicating that a single G deletion between bp1969-1976 of the RNF43 open reading frame indeed shift the frame and extend the length of the protein beyond G659. Subsequent work only used G659Vfs41 as this is the actual cancer mutation.

**Figure 2.**
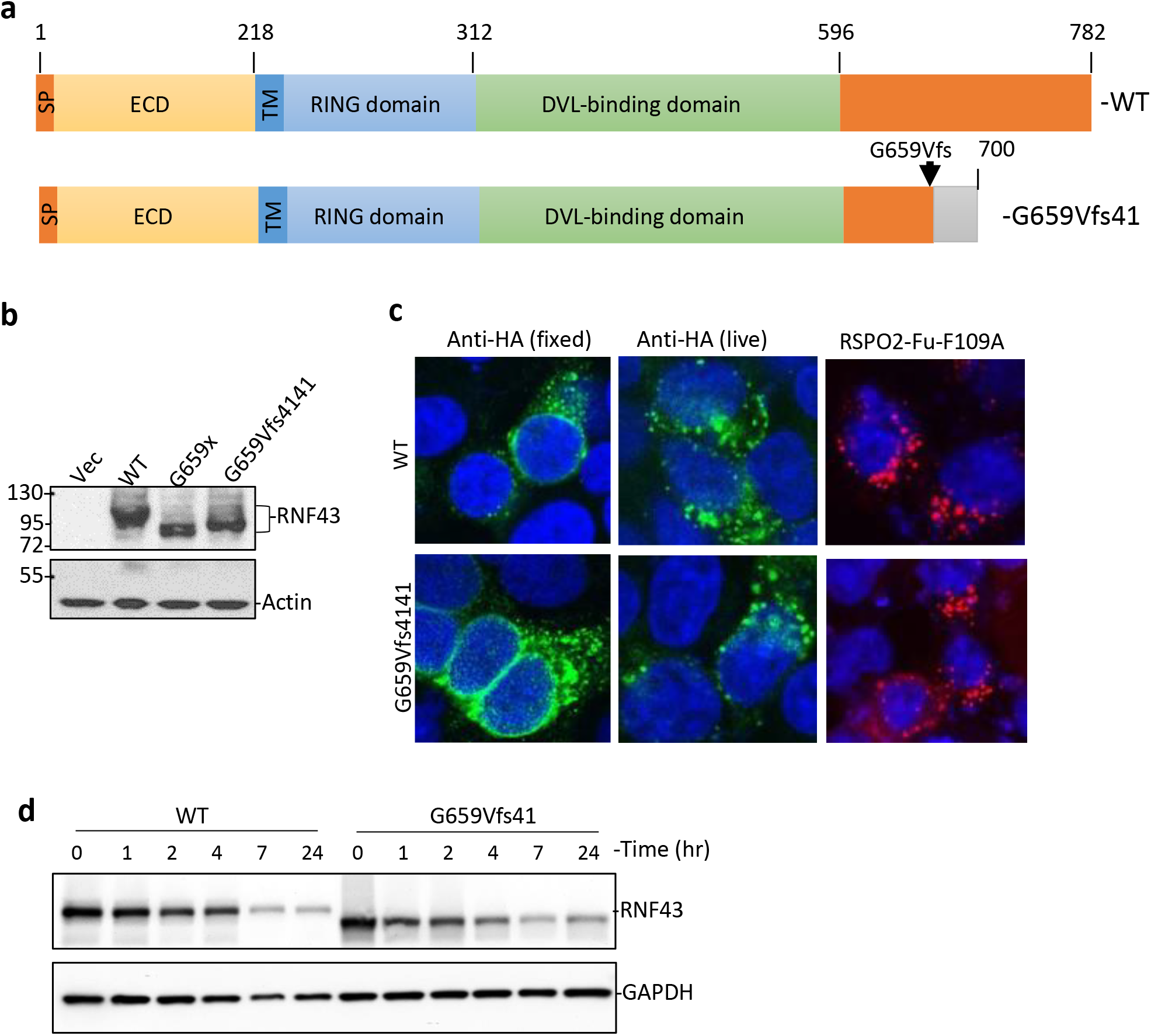
Expression, localization, RSPO binding, and stability of RNF43-G659Vfs41 is similar to RNF43-WT. a, Schematic diagram of domain structures of RNF43-WT and RNF43-G659Vfs4141. ECD, extracellular domain; TM, transmembrane domain; RING, RING E3 ligase domain. b, WB of HEK293T cells transfected with vector control (Vec), or HA-tagged RNF43-WT, -G659x, or -G659Vfs41 using anti-HA antibody. Actin is loading control. c, Confocal immunofluorescence of STF cells with double KO of RNF43 and ZNRF3 (D8) transfected with RNF43-WT and -G659Vfs41 using mouse anti-HA and Alexa488-labeled anti-mouse IgG (green). Nuclei were counter-stained with TO-PRO-3 Iodide (blue). d, stability of RNF43-WT and G659Vfs4141 in HEK283T cells following cycloheximide treatment.

To examine functions of the RNF43 mutant, we generated a double knockout of RNF43 and ZNRF3 in the Wnt/β-catenin signaling reporter cell line HEK293-STF to remove endogenous activity of the two E3 ligases using the CRISP/Cas9 method as previously described^5,23,24^. As expected, the double knockout cells (STF-RZ-DKO) had higher response to Wnt stimulation and no longer responded to RSPO1 treatment (Supplementary Fig. S1a). HA-tagged RNF43-WT and –G659Vfs41 were then transfected into STF-RZ-DKO cells and examined by immunofluorescence. In fixed and permeabilized cells, staining with anti-HA antibody showed that both WT and G659Vfs41 were mostly found in intracellular vesicles with strong perinuclear localization (Fig. 2c, left panel). In live cell staining, anti-HA antibody was rapidly internalized into vesicles in both WT and G659Vfs41-transfected cells (Fig. 2c, mid panel). To test if the mutant was able to bind to RSPO, we used the fusion protein RSPO2-Fu-F109A-IgG1-Fc which binds to ZNRF3/RNF43 with high affinity but not to LGR4 or LGR5^25^. In both WT- and G659Vfs41-transfected cells, RSPO2-Fu-F109A showed strong binding and rapid internalization (Fig. 2c, right panel). We then examined if the mutant had altered post-translational stability due to truncation and addition of the neopolypeptide. STF-RZ-DKO cells with transfected with either RNF43-WT or –G659Vfs41, and treated with cycloheximide. The level of both WT and G659Vfs41 decreased to a similar extent following the addition of cycloheximide (Fig. 2d), suggesting that two proteins had similar post-translational stability. Overall, these results indicate that RNF43-G659Vfs41 is nearly identical to WT in expression, localization, RSPO binding, and protein stability.

### RNF43-G659Vfs41 is similar to RNF43-WT in inhibiting Wnt/β-catenin signaling and decreasing Wnt receptor level

We first compared functional activity of RNF43-WT and -G659Vfs41 in the STF-RZ-DKO cells using the TOPFlash Wnt/β-catenin reporter enzyme assay^23^. Transfection of either WT or G659Vfs41 led to approximately 50% decrease in reporter enzyme activity (Fig. 3a), and RSPO2 was able to restore the activity to the level of vector control cells in a dose-dependent manner (Fig. 3a) even though the potency of RSPO2 was much lower than in regular STF cells^25^. WB analysis of the cells showed both WT and mutant were expressed at similar levels (Fig. 3b). The incomplete suppression of Wnt activity in the TOPFlash assay (Fig. 3a) was likely due to only approximately half of the cells expressing RNF43 following transient transfection based on immunofluorescence. The diminished potency of RSPO2 is likely an effect of high recombinant expression of RNF43 protein.

**Figure 3.**
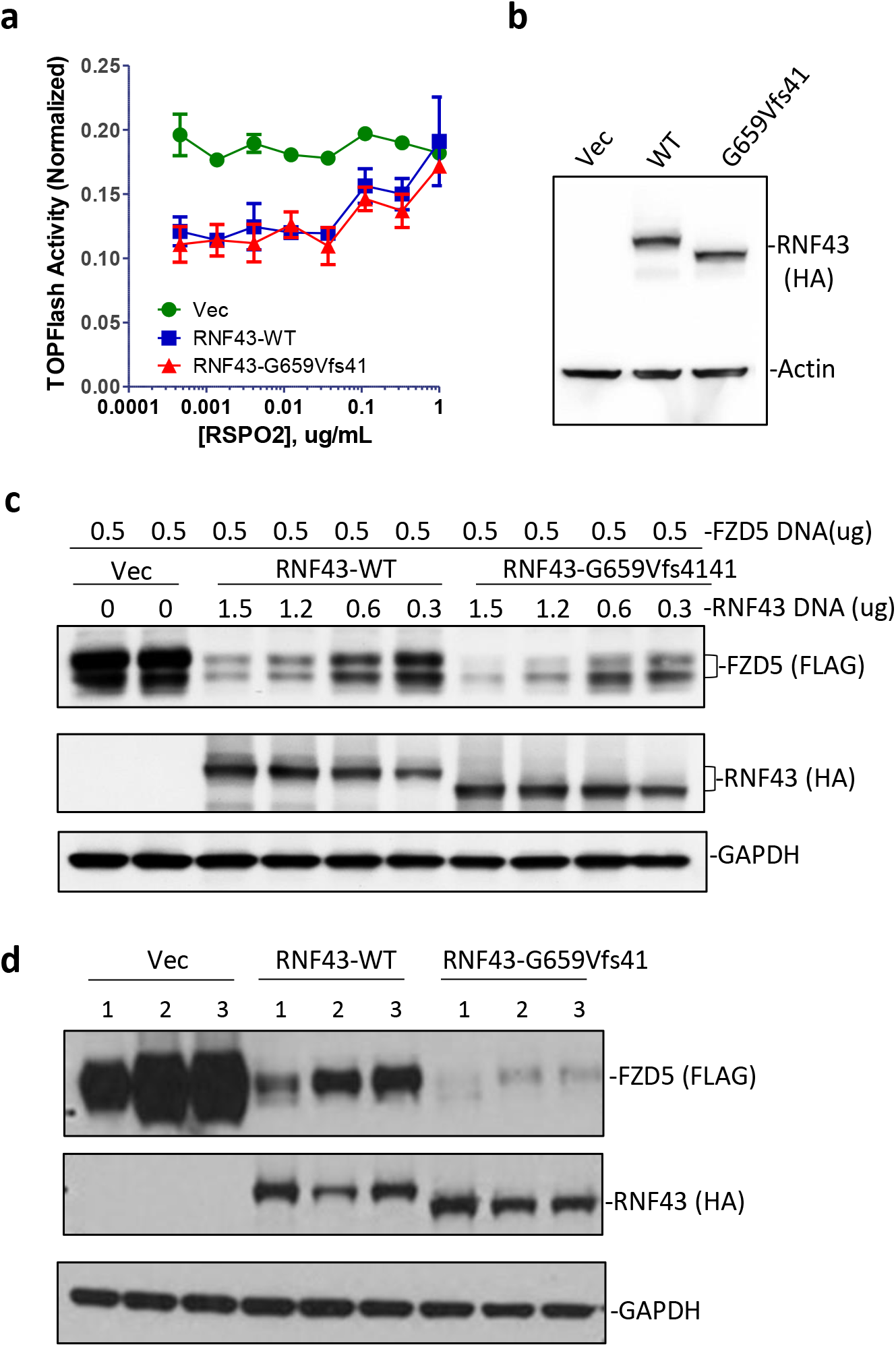
RNF43-G659Vfs41 inhibited Wnt signaling and reduced FZD levels as well as RNF43-WT. a, TOPFlash activity of STK-RZ-DKO cells transfected with vector (Vec), RNF43-WT, or –G659Vfs41 in response to RPSO2 treatment. b, WB of cells from a with anti-HA antibody. c, WB of FZD5 in STK-RZ-DKO cells transfected with Vector, RNF43-Wt and –G659Vfs41 with anti-FLAG and anti-HA antibody. d, WB of FZD5 of HEK293T cells transfected with vector, RNF43-WT or G659Vfs31 and treated with vehicle or RSPO2-Furin-Fc proteins (R2Fu) at 10 ug/ml. 1= vehicle; 2 = R2Fu-WT; 3 = R2Fu-F109A. Actin and GAPDP are loading control.

We then directly examined changes in the level of FZD protein following co-transfection of FLAG-tagged FZD5 and various amounts of WT or mutant RNF43 DNA into STF-RZ-DKO cells. WB analysis of FZD5 showed an approximate inverse correlation between levels of FZD5 and expression of either RNF43-WT or G659Vfs4141 (Fig. 3c), consistent with the expected function of RNF43 which is to ubiqutinate FZD for degradation^2,3^. Furthermore, RSPO2-Fu-WT or –F109A were able to increase levels of FZD in HEK293T cells transfected with either RNF43-WT or –G659Vfs41 to a similar extent (Fig. 3d), consistent with the binding and reporter enzyme activity data of WT and mutant RNF43. Intriguingly, in FZD degradation experiments, cells transfected with G659Vfs41 appeared to have lower levels of FZD5 when compared to those in WT-transfected cells (Fig. 3c-d), suggesting that the mutant might be even slightly more active. Overall, these functional results clearly indicate that RNF43-G659Vfs41 was fully capable of suppressing Wnt/β-catenin signaling and reducing FZD levels as well as RNF43-WT did.

### RNF43-G659Vfs41 is associated with low Wnt signaling activity and BRAF-V600E mutation only in colon cancer

RNF43-G659Vfs41 had been suggested to drive or facilitate colon cancer formation based on the presumption that it no longer functions in inhibiting Wnt signaling^7,9,14,26^. Since RNF43 is a target of Wnt signaling itself^2,3^, we examined the expression of RNF43 in relation to its own mutation status. Surprisingly, RNF43 expression in tumors with G659Vfs41 mutation was much lower than in those without any mutation (mean RNA-seq RSEM value of 1746 vs 6897, p < 0.05, Fig. 4a). Tumors with non-G659Vfs41 mutations in RNF43 also had lower expression of RNF43 (mean RNA-seq RSEM value = 3337, p < 0.05, Fig. 4a). However, in stomach and endometrial cancers, expression of RNF43 in G659Vfs41 or non-G659Vfs41 tumors was not different from those without any RNF43 mutation (Fig. 4b-c). Of note, expression of RNF43 in non-RNF43-mutated colon tumors was much higher than those in endometrial and stomach cancers (Fig. 4a-c). In fact, absolute mRNA levels of RNF43 in RNF43-G659Vfs41 colon cancer tumors is nearly identical to those in endometrial and stomach cancer. As stomach and endometrial cancers are rarely mutated in APC and colon tumors with RNF43 mutation are nearly exclusive with APC mutation, these data suggest that low expression of RNF43 mutations in colon tumors with RNF43 mutations was likely due to lack of APC mutations rather than due to RNF43 mutation itself. We then asked how ZNRF3 is related to RNF43 in expression and mutation status in colon cancer as both genes are targets as well as negative regulators of Wnt signaling. RNF43 and ZNRF3 expression are highly correlated (Spearman correlation efficient R = 0.622, Fig. 4d). Overall, these data suggest that tumors with RNF43 mutations had low expression of both RNF43 and ZNRF3, which implies low Wnt signaling even though low RNF43 and ZNRF3 tumors would have been previously predicted to have higher Wnt receptors levels and stronger Wnt signaling.

**Figure 4.**
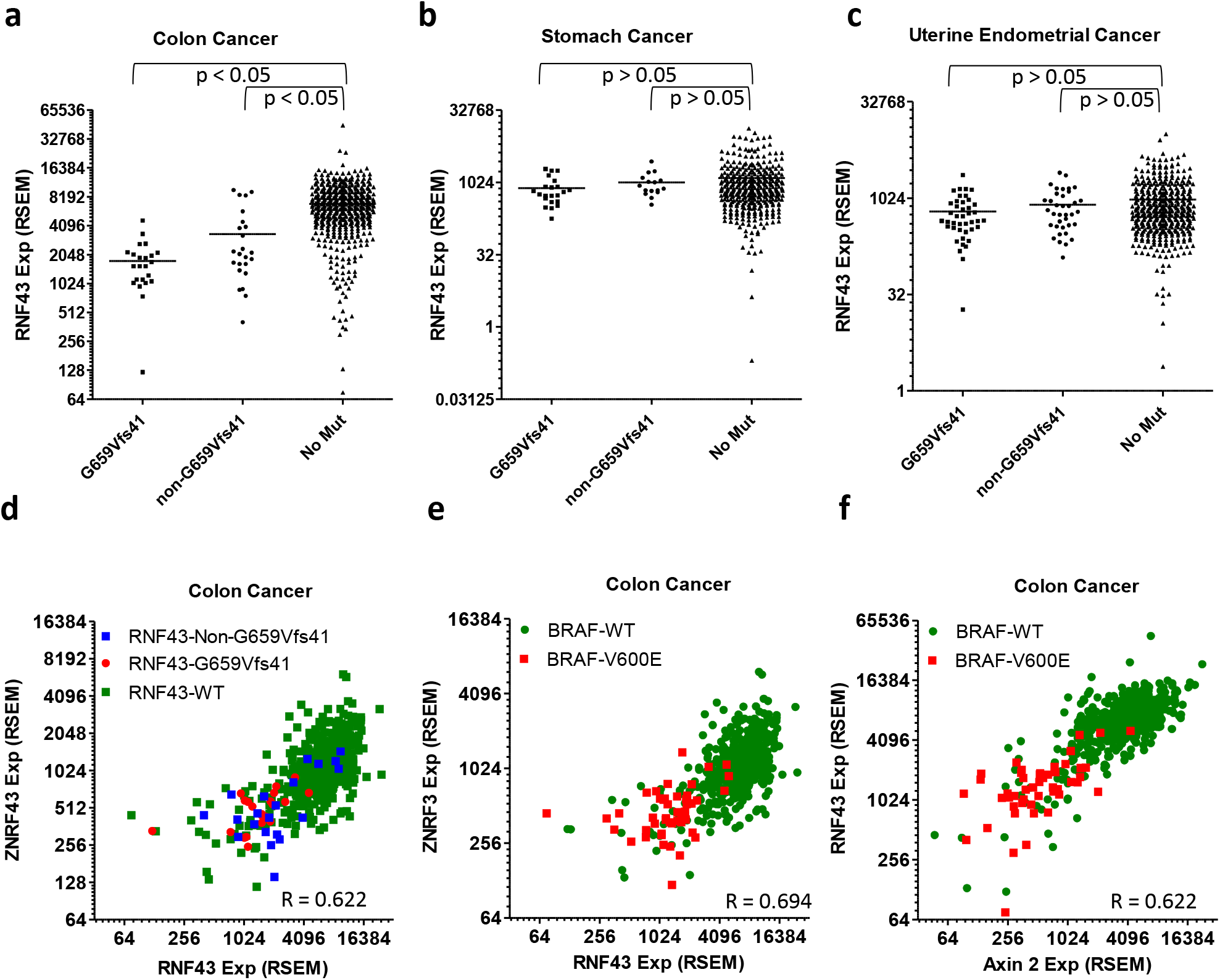
Expression of Axin2, RNF43, and ZNRF43 are highly correlated with each other and is low in RNF43-G659Vfs41 and BRAF-V600E tumors only in colon cancer. a, Scatter plot of expression level of RNF43 vs its own mutation status in colon cancer. b, Expression level of RNF43 vs its own status in stomach cancer. c, Expression level of RNF43 vs its own status in endometrial cancer. d, correlation between RNF43 and ZNRF3 in expression in relation to the mutation status of RNF43. e, correlation between RNF43 and Axin2 expression in relation to BRAF-V600E. f, correlation between RNF43 and ZNRF3 expression in relation to BRAF-V600E.

Previously, it was also noted that RNF43-G659Vfs41 is frequently associated with BRAF-V600E mutation^9,14,26^, an expected correlation since BRAF-V600E is most frequent in MSI-H tumors with low MLH1 expression^27^. In the TCGA colon cancer cohort with 533 sequenced tumors, there are 21 tumors with G659Vfs41 mutations and 47 tumors with BRAF-V600E mutation. Yet 14 of the 21 tumors with G659Vfs41 mutation also had BRAF-V600E mutation (p = 0.0001, Fisher’s exact test), indicative of a strong association between the two mutations. As colon tumors with MSI-H and BRAF-V600E mutations are also known to have low Wnt signaling^27^, we analyzed expression levels of Axin2, a prototypical Wnt signaling target, and ZNRF3, relative to RNF43 and BRAF mutation status in the colon cancer cohort. Axin2 and ZNRF3 mRNA levels are highly correlated with that of RNF43 across all samples (overall Spearman’s correlation coefficient = 0.694 for Axin2, 0.622 for ZNRF3, Fig. 4e-f), consistent with all three being Wnt target genes in the intestinal tract. Remarkably, nearly all BRAF-V600E tumors had low expression of Axin2, RNF43, and ZNRF (Fig. 4e-f), but not vice versa, consistent with previous report of low Wnt signaling in colon tumors with MSI-H BRAF-V600E mutation^27^. Overall, these genomic data clearly indicate that a subset of colon cancers, including nearly of all those with BRAF-V600E mutation, had low Wnt signaling as evident from their low expression of Axin2, ZNRF3, and RNF43

### Colon cancer cell line with RNF43-G659Vfs41 and BRAF-V600E mutation is sensitive to activation of Wnt signaling

As the majority of RNF43-G659Vfs41 mutations co-existed with BRAF-V600E and had low Wnt signaling, including low RNF43 and Axin2 expression, we mined the CCLE database for cell lines with these characteristics. The cell line RKO had the RNF43-G659Vfs41 mutation as confirmed by Bond et al^9^, BRAF-V600E and low Axin2 expression without APC mutation, an apparent representative of the most common RNF43-G659Vfs41 primary tumors. We also identified three other cancer cell lines with differential profiles in Wnt signaling gene expression and mutations as listed in Supplementary Table S1. Sensitivities of the four cell lines to various inhibitors in cell growth were then compared side-by-side: BRAF inhibitor PLX4720^28^, porcupine inhibitor LGK974 (inhibition of Wnt ligand secretion)^17^, tankyrase inhibitor XAV939 (stabilization of Axin1/2 and thus degradation of β-catenin)^29^, and GSK3 inhibitor CHIR99021 (inhibition of β-catenin phosphorylation and thus stabilization of β-catenin)^30^. Inhibition of BRAF (PLX4720) was highly effective in HT29 cells (BRAF-V600E, RNF43-WT, APC mutant) but had little activity in RKO cells (BRAF-V600E, RNF43-G659Vfs41, APC-WT) (Fig. 5a), consistent with previously published results by others^31^. In LoVo and DLD1 cells (no RNF43 nor BRAF mutation), PLX4720 had intermediate effect (Fig. 5a). Remarkably, activation of Wnt signaling by the GSK3 inhibitor CHIR99021 was able to inhibit the growth of RKO cells completely with only minor effect on the other three cell lines at 10 uM (Fig. 5b). On the other hand, inhibition of Wnt signaling by either blockade of Wnt ligand synthesis (LGK974) or stabilization of Axin1 (XAV939) had little effect on the four cell lines (Fig. 5c-d) despite APC mutation in all cell lines except RKO. These data suggest that activation of Wnt signaling via inhibition of GSK3 is detrimental to RKO cells which have both RNF43-G659Vfs41 and BRAF-V600E, suggesting that BRAF-V600E mutation is not compatible with high Wnt signaling under certain circumstances. This may be the underlying reason of why only MSI-H BRAF-V600E tumors had low Wnt signaling whereas BRAF-V600E tumors without MSI-H had high Wnt signaling^27^. Of note, HT29 cells are MSS (microsatellite stable) whereas RKO cells are MSI-H^32^.

**Figure 5.**
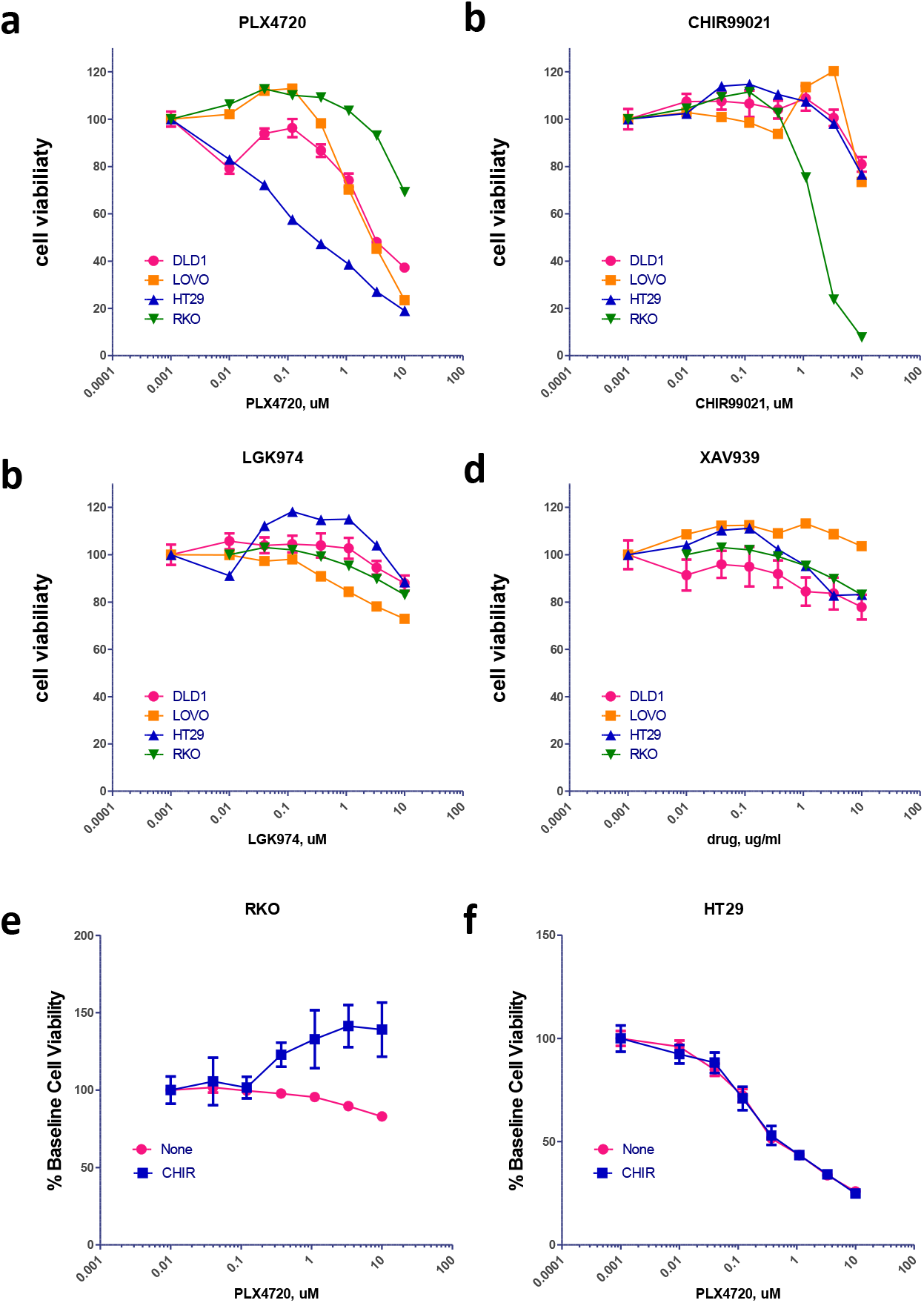
Cell viability of colon cancer cell line in response to treatment with inhibitors and activators of Wnt and BRAF signaling. a, BRAF inhibitor BRAF-V600E. b, Wnt/β-catenin signaling activator CHIR99021. c, Wnt ligand secretion inhibitor LGK974. d, Wnt/β-catenin signaling inhibitor XAV939. e-f, Combined effect of CHIR and PLX4720 in RKO (e) and HT29 cells (f). For e and f, varying concentrations of PLX4720 was tested together with vehicle or CHIR99021 at a fixed concentration of 3 uM.

If high Wnt signaling due to GSK3 inhibition is incompatible with high BRAF activity (BRAF-V600E), we would expect that inhibition of BRAF may rescue the cells from high Wnt signaling. To test this, we compared the effect of combining PLX4720 with CHIR99021 in RKO and HT29 cells. In RKO cells, PLX4720 had little effect by itself but was able to partially rescue the effect of CHIR99021 which inhibited cell growth by ~50% at the used concentration (3 uM) (Fig. 5e). In HT29 cells, CHIR99021 at the same concentration had no effect on PLX4720 (Fig. 5f). These data strongly suggest that in RKO cells, cell growth inhibition by CHIR99021 depended on high BRAF activity, consistent with the model that high Wnt signaling is not tolerated with BRAF-V600E in MSI-H tumors.

### GSK3 inhibition in RKO cells led to stabilization of β-catenin

To determine the mechanism of sensitivity of RKO cells to GSK3 inhibition, we first examined Wnt and MAPK signaling pathways in the four cell lines. RKO cells had little β-catenin in the cytoplasm at base line and CHIR99021 treatment increased its level as expected (Fig. 6a). CHIR99021 also increased the level of cytoplasmic β-catenin in HT29, and LoVo cells but not in DLD1 cells (Fig. 5A). Level of Axin2 was only increased in RKO cells with CHIR99021, probably because the other three cell lines are mutated in APC and thus had already saturated in Axin2 expression. Levels of Axin1 were decreased in all CHIR99021-treated cell lines except RKO, likely due to degradation of the β-catenin destruction complex as a result of lack of phosphorylated β-catenin. In contrast, PLX4720 had no effect on the markers of Wnt signaling pathway in the four cell lines (Fig. 6a). In the MAPK pathway, HT29 cells showed the best response in pERK reduction with PLX4720 treatment, consistent with its high sensitivity. RKO cells, despite having BRAF-V600E mutation, only had a minor reduction in pERK. LoVo cells showed an increase in pERK due to positive feedback following inhibition of BRAF in cells with KRAS mutation. We then asked if high Wnt signaling would induce apoptosis in RKO cells. As shown in Fig. 6c, a significant increase of PARP cleavage was observed in RKO cells treated with CHIR99021 but not PLX4720. Overall, these results confirmed that GSK3 inhibition in RKO cells resulted in dramatic increase in the level of cytoplasmic β-catenin and Axin2, which led to apoptosis and cell death.

**Figure 6.**
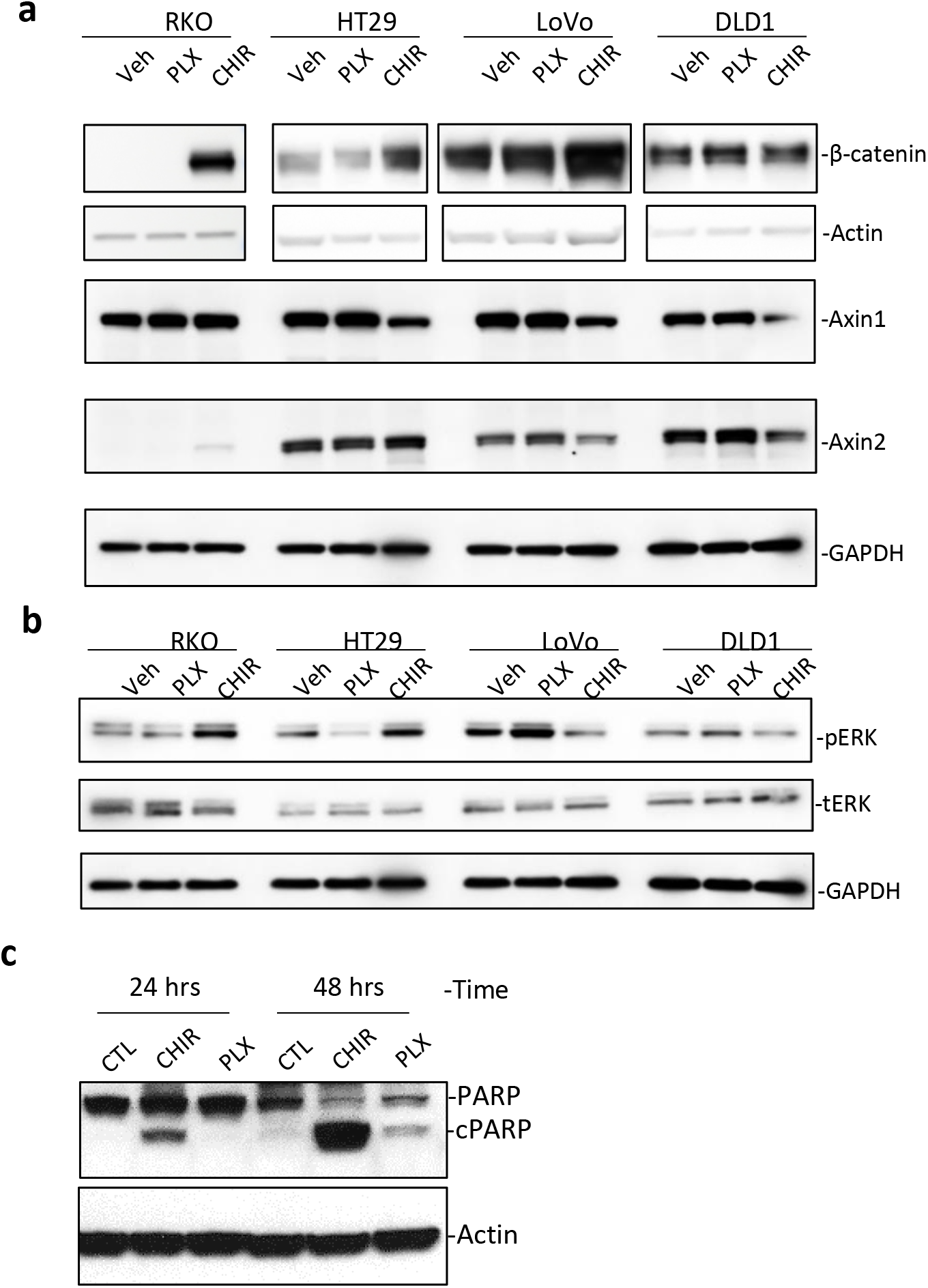
CHIR99021 induced apoptosis in RKO cells by stabilizing β-catenin. a, WB analysis of Wnt signaling pathway indicators in the four cell lines treated with vehicle (CTL), CHIR99021 (CHIR), or PLX4720, each at 3 uM for 24 hours. b, WB analysis of phosphor-ERK (pERK) and total ERK (tERK) in the four cell lines treated with the indicated compound at 3 uM for 24 hours. c, WB of PARP1 cleavage in RKO cells following treatment with CHIR or PLX at 3 uM for the indicated period of time. cPARP1 = cleaved PARP1. Actin and GAPDH were used as loading control.

## Discussion

RNF43 and its related homolog ZNRF3 are key negative regulators of Wnt signaling and both were found to be mutated in all major types of solid tumors at relatively low frequency. Some of the null or missense mutations of RNF43 located in the N-terminal half had been shown to be loss-of-function and presumably play a role in tumorigenesis of the pertinent tumors^4,17^. The most common recurrent mutation is RNF43-G659Vfs41 which has been “automatically” assumed to be a loss-of-function mutation that would lead to increase in Wnt signaling and thus drive or facilitate tumorigenesis. Yet, no evidence has been presented to show that the mutation abolishes the enzyme function as a driving mechanism in oncogenesis. Here we show that this mutation does not alter the function of RNF43 in suppressing Wnt signaling, and colon tumors with RNF43-G659Vfs41 mutation had low Wnt signaling and may even be sensitive to activators of Wnt signaling.

For both ZNRF3 and RNF43, three domain (ECD, RING E3 ligase domain, and DVL-binding domains) were shown to be essential for their activity in ubiquitinating FZD and inhibiting Wnt signaling^4,5^. RNF43-G659Vfs41 replaces the C-terminal 124-AA region of RNF43 with a 41-AA neopeptide that shows no homology to any other protein or functional motif. The replaced region is far downstream of the DVL-binding domain and previous studies by others showed that deletion of the entire region downstream AA-596 (end of DVL-binding domain) had no effect on RNF43’s activity in inhibiting Wnt signaling^4,5^. Complete deletion of the last 124-AA was not expected to have any effect. Yet, this mutation has been proposed to be a driving mechanism of oncogenesis in the colon, largely because it is a frameshift/truncation mutation of a strong negative regulator of Wnt signaling has near exclusivity with APC mutation. In theory, it is possible that the 41-AA neopolypeptide could somehow inhibit the function of the enzyme and increase Wnt signaling. Here we excluded this possibility by demonstrating that the expression, localization, and function of recombinant RNF43 with the very G659Vfs41 mutation were no different from RNF43-WT. In fact, the mutant appeared to be slightly more active in some of the experiments. In cancer cells, the mutant RNF43 would co-express with WT RNF43 and ZNRF3, and could possibly inhibit the function of the WT enzymes if WT and mutant enzymes could form inactive dimers. However, RNF43 was shown to not to form dimers in crystal structure^33^, and transfection of RNF43-G659Vfs41 into HEK293T cells with endogenous expression of ZNRF3 still suppressed caused degradation of FZD (Fig. 3d). Therefore, RNF43-G659Vfs41 is not expected to affect enzyme activity of WT RNF43 and ZNRF3 in cancer cells that co-express these three forms of the E3 ligases to increase Wnt signaling, and thus the mutant is unlikely to play a driving role in the formation or progression of cancers with this mutation.

Multiple studies reported the association of RNF43-G659Vfs41 with BRAF-V600E in MSI-H colon tumors ^9,12–14,26^. However, the allele frequency of RNF43 and BRAF mutations were not correlated (supplementary Fig. S3), suggesting that the events were independent. Also, in stomach and endometrial cancers in which BRAF-V600E was not found, RNF43-G659Vfs41 occurred in a frequency similar to those of MSI-H colon cancer. In all three cancers types with high incidence of RNF43-G659Vfs41, tumors with G659Vfs41 all had low expression of MLH1 and presumably defective DNA repair, suggesting that replication of the seven G repeat in coding sequence of RNF43 is very prone to error and highly dependent on MLH1 for correction. Together with the finding that G659Vfs41 had no effect on RNF43’s activity in Wnt signaling, these data strongly suggest that G659Vfs41 is a passenger mutation that happens to occur at high frequency due to the lack of MLH1. Interestingly, Roudko et al just reported that RNF43-G659Vfs41 was expressed as a neopolypeptide that is predicted to be a highly immunogenic tumor antigen in the MSI-H subtypes of the three tumor types (colon, stomach, and endometrial cancers)^34^.

The association of BRAF-V600E and low Wnt signaling was only found in MSI-H colon tumors which are rarely mutated in APC^27^. In melanoma cells with BRAF-V600E mutation, inhibition of BRAF by PLX4720 led to cell death and was dependent on increased Wnt/β-catenin signaling as a result of decreased Axin1, a negative regulator of β-catenin^35^. In colon cancer cell lines with or without BRAF-V600E mutation, we found that PLX had no effect on the level of Axin1 nor β-catenin (Fig. 6A), indicating that BRAF-V600E had distinct effect in colon cancer and melanoma cells. In cell growth tests of the two colon cancer cell lines with BRAF-V600E mutation, HT29 (MSI-low, not hypermutated) and RKO (MSI-HI)^32^, HT29 cells were highly sensitive to PLX4720 whereas RKO cells were rather resistant which is consistent with previous report^31^. Instead, we found RKO cells is highly sensitive to a Wnt/β-catenin signaling activator CHIR-99021 which was not known to be cytotoxic to any cell line. In melanoma cells with BRAF-V600E mutation, inhibition of BRAF by PLX is synergistic with activation of Wnt signaling in causing cell death^35^, suggesting that low BRAF activity is incompatible with high Wnt signaling. In RKO cells, however, PLX4720 would slightly counteract the effect of CHIR99021, suggesting that low BRAF activity is more compatible with high Wnt signaling. This would imply that high BRAF activity in BRAF-V600E cells would require low Wnt signaling, consistent with low levels of Axin2 and other Wnt target genes as well as the lack of APC mutation in MSI-H colon cancer cells with BRAF-V600E mutation^36^. Interestingly, in intestinal stem cells, high Wnt signaling and MAPK activity were shown to be incompatible as the two pathways seem to inhibit each other^35,37,38^. Taken together, these findings suggest that MSI-H tumors with BRAF-V600E mutation had low Wnt signaling due to inhibition the hyperactive MAPK signaling. More importantly, colon cancers of MSI-H with BRAF-V600E tumors may not tolerate high Wnt signaling and activation of Wnt signaling in such tumors may provide a therapeutic approach.

In conclusion, we found that RNF43-G659Vfs41wa able to inhibit Wnt signaling as well as the WT enzyme did and that this mutation is most likely a passenger mutation due to error-prone replication of a seven G repeat in its open reading frame in MLH1-deficient tumor cells. Its coexistence with BRAF-V600E is a pure coincidence for both being enriched in MSI-H tumors. Colon cancer cells with RNF43-G659Vfs41 and BRAF-V600E mutations had low levels of Wnt signaling and are not sensitive to inhibitors of Wnt signaling. To the contrary, such tumors may respond to the treatment by Wnt activators.

## Materials and Methods

### Plasmids and Cloning

Plasmids encoding N-terminal HA-tagged RNF43-WT and -G659x with a CD8 signal peptide were cloned into pIRESpuro3 by standard PCR and In-Fusion HD cloning kit (Clonetech). HA-RNF43-G659Vfs41 was then generated by deleting a G in bp 1979-1976 of the RNF43 open reading frame in the HA-RNF43-WT plasmid by site-directed mutagenesis. FLAG-tagged FZD5 was cloned into pIRESpuro3 as described previously^25^. All plasmids were verified by complete DNA sequencing.

### Recombinant proteins, antibodies, and Western blotting

Recombinant full-length human RSPO2 was purchased from R&D Systems. RSPO2-Fu-WT and RSPO2-Fu-F109A were purified as described previously ^25^. For western blot analysis, anti-HA (Invitrogen cat #71-5500), anti-FLAG (Sigma cat # F7425), anti-β-actin (Cell Signaling cat #4970), anti-β-catenin (Cell Signaling cat #9562), anti-Axin1 (Cell Signaling cat #2087), anti-Axin2 (Cell Signaling cat #2151), anti-phospho-ERK (Cell Signaling cat #9101), total ERK (Cell Signaling cat #4695), anti-PARP (cell signaling cat #9532), and anti-GAPDH (Cell Signaling cat #2118) were used. For ICC, anti-HA-Alexa 488 (Cell Signaling cat #2350), anti-human-Alexa 488 or –Alexa 555 (Invitrogen cat #A11013 and #A21433) were used. For western blotting, cells were lysed with RIPA buffer (50 mM Tris-Cl pH 7.4, 150 mM NaCl, 1mM DTT, 1 % Triton X-100, 1 % sodium deoxycholate, 0.1 % SDS), supplemented with protease and phosphatase inhibitors, and reduced at 37 °C for 1 hr. HRP-conjugated secondary rabbit or mouse antibodies (Cell Signaling) were used following the standard ECL protocol.

### Cell culture, transient transfection, and protein stability analysis

All cell lines were purchased from the ATCC and cultured in a 37 °C humidified incubator containing 5% CO2. HEK293T, HEK293-STF and HT29 cells were cultured in DMEM, DLD1 and Lovo cells in RPMI-1640, and RKO cells in MEM media. All media were supplemented with 10% FBS, 100 units/mL penicillin, and 100 mg/mL streptomycin. For transient transfection, ~80 % confluent cells were transfected with DNA: FuGENE HD (Promega) ratio of 1:3 for all transient transfection presented. For the protein stability analysis, cells with treated with cycloheximide at 100 ug/ml for the indicated periods of time and harvested, followed by WB analysis.

### Generation of STF RNF43 and ZNRF3 double knockout cells

For CRISPR-based knockout of ZNRF and RNF43 in HEK293-STF cells, the guide sequences GCAGGGTAGCCATCAGCAGCC (corresponding to nucleotide 44-64 of human RNF43 open reading frame) and AGGACTTGTATGAATATGGC of ZNRF3 (corresponding to nucleotide 338-357 of human ZNRF3 open reading frame) were cloned into the vector Lenti-CRISPR2 as described^24^. The two guide sequences were provided by Dr. Feng Cong as they were used in the generation of double knockout of RNF43 and ZNRF in HEK293 cells^5^. Lentiviral particles of lenti-CRISPR2 containing RNF43 and ZNRF3 guide sequence were used to co-infect HEK293-STF cells and cells were selected with puromycin at 1 µg/ml. Single colonies were isolated and analyzed for Wnt and RSPO response by the TOPFlash assay^23^. One clone (D8) showed complete loss of response to RSPO1 in the TOPFlash assay (supplementary Fig. S1a) and knockout of both RNF43 and ZNRF3 in D8 cells were verified by sequencing the genomic regions flanking the two guide sequences. In brief, genomic sequences were amplified by PCR using forward primers CGAAGTGACATTCAATCACAAG and GACGTTACTTTTGGCTATAGCATCTG (located in intron 1 of RNF43 and ZNRF3, respectively) and reverse primers CCTTCTGCTGGAGTTATTTCAGC and CGACAAGAGGGTAGAGCCCGCTC (located in intron 2 of RNF43 and ZNRF3, respectively). The PCR products were cloned into pCR2.1 vector using TA cloning kit (Invitrogen) and a total of 16 clones were sequenced with the identified mutations shown in Supplementary Figure S1b.

### Wnt/β-catenin signaling TOPFlash reporter enzyme assay and immunofluorescence

TOPFlash assay in STF RNF43-ZNRF3 double KO (RZ-DKO) cells transfected with vector, RNF43-WT, or RNF43-G659Vfs41 were carried out as described previously^25^. All TOPFlash experiments were repeated at least three times with duplicates or triplicates in each experiment. For RNF43 localization, STF-RZ-DKO cells transfected with RNF43-WT or G659Vfs41 were treated as indicated. The treated cells were incubated at 37 °C with 95 % humidity and 5 % CO_2_ for 1 hr and fixed with 4.2 % paraformaldehyde, followed by permeabilization with 0.1 % saponin. Then, the secondary antibody, anti-human-Alexa 555 was used to label Fc-tagged RSPO2. Cells were imaged under the confocal microscope and analyzed by the Leica LAS AF Lite software.

### In vitro cell viability assays

Cells were seeded at various numbers (depending on growth rate) in half-area white 96-well plates and serial dilutions of the compounds were added. The cells were incubated for 4 days and cell viabilty was measured using CellTiter-Glo^®^ Luminescent Cell Viability Assay (Promega).

### Cancer genomics data and statistical analysis

A ll expression and mutation data of TCGA tumor samples were doanloaded from cBioportal^20,21^. Data plotting and statistical analysis were carried out using GrpahPad Prism.

## Acknowledgement

We would like to thank cBioPortal for the support of the database and analysis tools

## Funding

Cancer Prevention and Research Institute of Texas (RP160235 and RP170245 to Q.J.L.), National Institute of Health (R01GM102485 to Q.J.L and R01CA226894 to K.S.C.), and Gordon Endowment for Bowel Cancer Research (to Q.J.L.).

## Competing interests

The authors declare no competing interests.

## Author Contributions

Q.J.L. analyzed data and wrote the manuscript. J.T., S.P., W.Y., S. Z., and W.L. and K.S.C. performed the experiments analyzed data. S.P. and K.S.C. edited the manuscript.

Supplementary information is available for this paper.

