## supplementary table and figures for "The most common RNF43 mutant G659Vfs41 is fully functional in inhibiting Wnt signaling and unlikely to play a role in tumorigenesis"

### Supplementary Tables and Figures

Supplementary Table S1. Expression and mutation profile of 4 colon cancer cell lines

| Cell line | Expression | | | Mutation | | | | |
|---|---|---|---|---|---|---|---|---|
| | AXIN2 | RNF43 | ZNRF3 | RNF43 | APC | BRAF | KRAS | CTNNB1 |
| HT29 | 6.3 | 9.9 | 10.3 | None | T1556Nfs*3, E853* | V600E,T119S | None | None |
| RKO | 4.7 | 5.0 | 6.6 | X659fs-hom | None | V600E | None | None |
| DLD1 | 8.6 | 9.6 | 9.8 | X659fs , L214M | S1426x | None | None | None |
| LOVO | 7.3 | 7.6 | 8.5 | None | R1114*, R2816Q, M1431Cfs*42 | None | G13D | None |

**a**

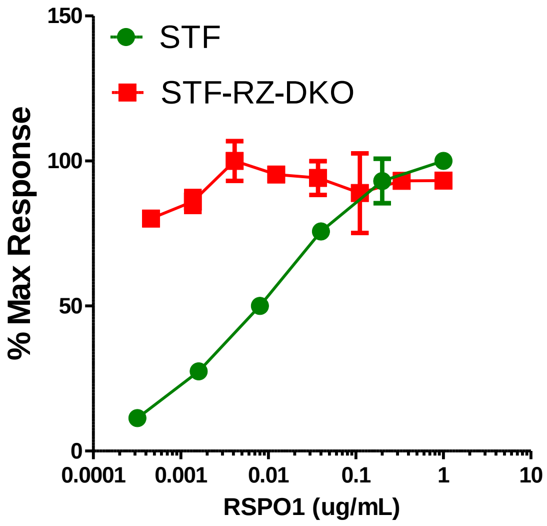


**b**

| | Sequences of ZNRF3 Exon 2 and RNF43 Exon 2 in the STF clone D8 that had knockout of both RNF43 and ZNRF3. | # nucleotides changed |
|---|---|---|
| ZNRF3 | 5′- CGAAGAGGACTTGTATGAATATGGCTGGGTAGGAGTGGTGAAGCT -3′ | |
| D8-1 | 5′- CGAAGAGGACTTGTATGAATAT**T**GGCTGGGTAGGAGTGGTGAAGCT -3′ | +1 |
| D8-2 | 5′- CGAAGAGGACTTG---------GGCTGGGTAGGAGTGGTGAAGCT -3′ | -9 |
| RNF43 | 5′- TGCCTGCAGGGTAGCCATCAGCAGCCAGGGCCAGAGGGCAGCCAGCTG -3′ | |
| D8-1 | 5′- TGCCTGCA-GGTAGCCATCAGCAGCCAGGGCCAGAGGGCAGCCAGCTG -3′ | -1 |
| D8-2 | 5′- TGCC----------------------------AGAGGGCAGCCAGCTG -3′ | -28 |

Supplementary Figure S1. Characterization of STF cells with double knockout of RNF43 and ZNRF3 (STF-RZ-DKO). a, TOPFlash Wnt signaling assay results of parental and RZ-DKO cells in response to RSPO1. b, Sequence alignment of exon2 of RNF43 and ZNRF3 in STF cells with double knockout of the two genes. For both genes, the sequences of each allele are aligned with that of the WT sequence with the deleted sequences shown as gaps.

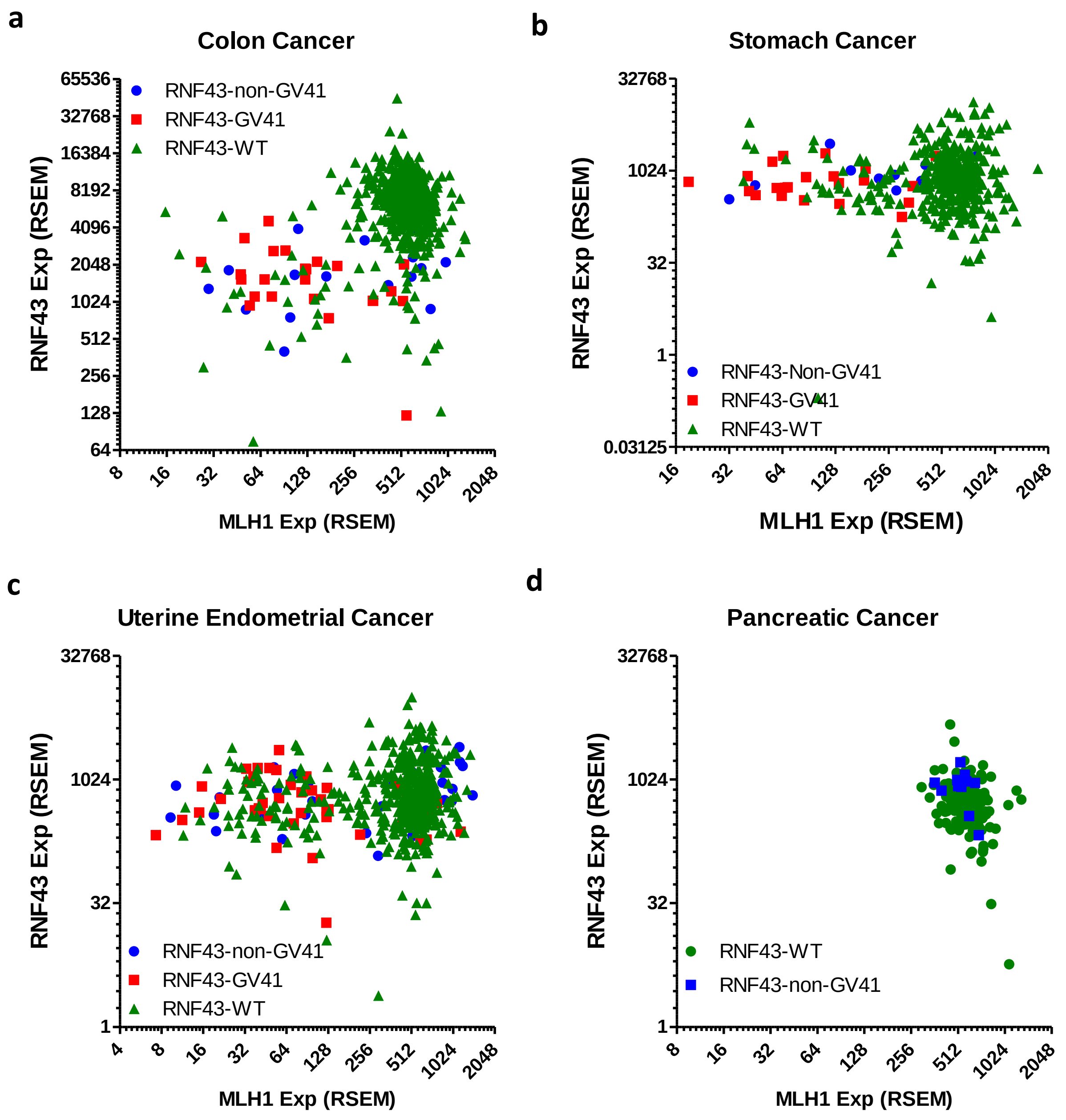


Supplementary Figure S2. RNF43-G659Vfs41 mutations in colon, stomach, and endometrial cancers occurred nearly exclusively in tumors with low MLH1 expression. a-d, RNA-seq expression data and mutation status of RNF43 were plotted against RNA-seq data of MLH1 in colon (a), stomach (b), uterine endometrial (c), and pancreatic cancer (d). GV41 = G659Vfs41.

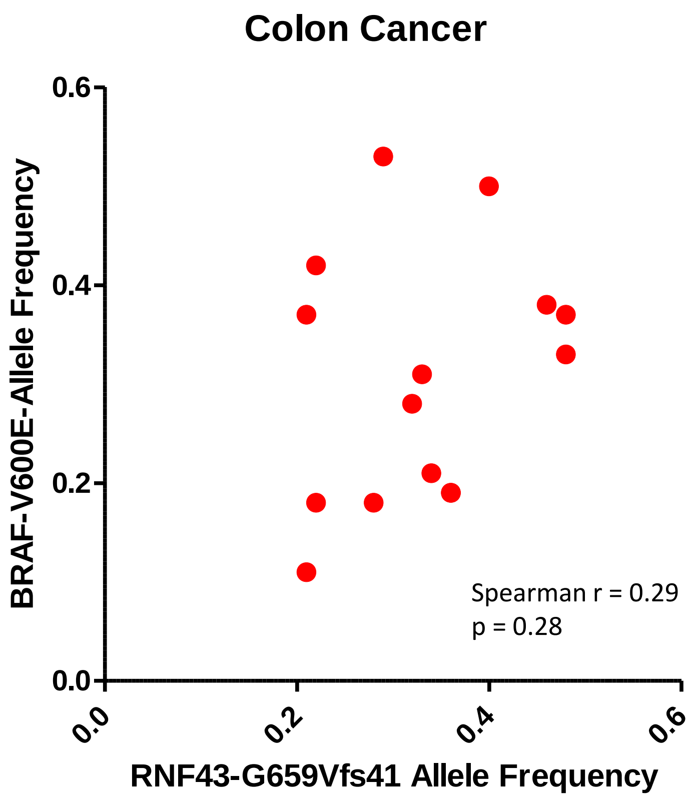


Supplementary Figure S3. No correlation between allele frequency of RNF43-G659Vfs41 and BRAF-V600E in TCGA's colon cancer cohort. Of the 15 tumors with both RNF43-G659Vfs41 and BRAF-V600E mutations, the respective allele frequencies were plotted as labeled.
